# PYEAST – Python Enabled Automated Strain Transformation

**DOI:** 10.1101/2025.05.19.655004

**Authors:** Abubakar Madika, Ankita Suri, Anjali Purohit, Damian Van Raad, Michael Norman, Carol J. Hartley, Thomas Loan

## Abstract

*Saccharomyces cerevisiae* is a widely used biotechnological workhorse in both academic and industrial settings. One reason for its continued popularity is the extensive legacy of genetic tools, developed over its long history of use, that enable precise manipulation of the *S. cerevisiae* genome. These tools have enabled extensive genetic characterisation and dramatic re-programming efforts for applications ranging from fundamental research to industrial chemical production. Here we present a digital toolkit called PYEAST (<u>Py</u>thon <u>E</u>nabled <u>A</u>utomated <u>S</u>train <u>T</u>ransformation) that encodes some of the most widely used methods for working with *S. cerevisiae* and modernizes them to leverage advances in DNA synthesis.

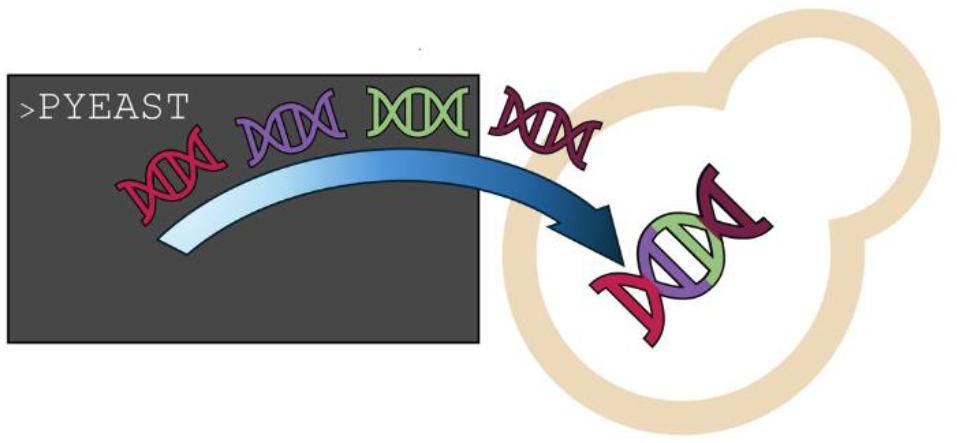

## Introduction

*S. cerevisiae* has a high propensity for homologous recombination (HR) over other DNA repair mechanisms.^1^ This preference for HR has been exploited in the development of a plethora of tools for genetic manipulation. These tools include assembly of large constructs,^2,3^ integration of new DNA sequences into chromosomal DNA,^4^ and scarless-deletions or replacements of genomic sequences.^5,6^ These tools have facilitated the construction of large neochromosomes^7^ and even entire genomes .^8,9^ Recently, CRISPR/Cas tools have become widely used,^10^ although outside of academic research the widespread use of CRISPR is still hampered by a complex intellectual property landscape.^11,12^

In establishing our own workflows for yeast synthetic biology, we encountered decades of improvements and optimizations on related techniques, including a series of sophisticated programs and tools for computer aided design.^13–15^ These efforts largely focused on plasmid assembly, and the use of automation tools in large-scale pipelines making them poorly suited to our specific needs. Like many laboratories, our work involved mostly low-to-medium throughput and required a wide variety of techniques for assembling constructs and modifying the *S. cerevisiae* genome. To accommodate these diverse workflows, while also facilitating standardization, protocol sharing, computer aided design and ongoing process improvements, we began encoding our work in python scripts. These initially basic scripts steadily evolved into a unified program called PYEAST (<u>Py</u>thon <u>e</u>nabled <u>a</u>utomatic <u>s</u>train transformation). PYEAST streamlines the management of DNA components and primers while remaining simple enough to be maintained with limited programming experience and easily integrated with existing DNA sequence and primer collections. We share it here in the hope that these tools will be useful for other researchers in a similar position and serve as a reference for groups seeking to develop their own pipelines and codebases.

PYEAST is a command line interface containing four separate functions that span **t**ransformation **<u>a</u>**ssisted **r**ecombination (***tar***) for plasmid assembly, ***integrate*** for inserting DNA into the genome, ***delete*** for scarless deletions and ***replace*** for scarless replacements of genome sequences. We also provide a range of useful sequences, many of which can be amplified directly from the yeast genome or from common plasmids, allowing others to begin using this tool kit without the need to first acquire a large collection of DNA components. All commands rely on a shared file structure for managing DNA sequences, primers, and experimental documentation, with standardized inputs and outputs that facilitate protocol sharing and reproducibility. Below, we describe each command’s functionality and use cases of each command. Detailed examples of their use are provided in the supplementary information.

### Tar and integrate

The first two commands, *tar* and *integrate*, follow a comparable workflow (figure 1). The *tar* command is designed for the assembly of plasmids by homologous recombination of PCR products upon transformation into *S. cerevisiae*.^2^ This allows for the assembly of complex constructs, without the necessity to consider sequence elements like the type II restriction sites required for golden gate assemblies. This is particularly useful for promoters and terminators, which often contain sequence elements that can make synthesis challenging or even impossible, such as long stretches of A or T. In many cases, these sequences can be directly amplified from genomic DNA.

**Figure 1:**
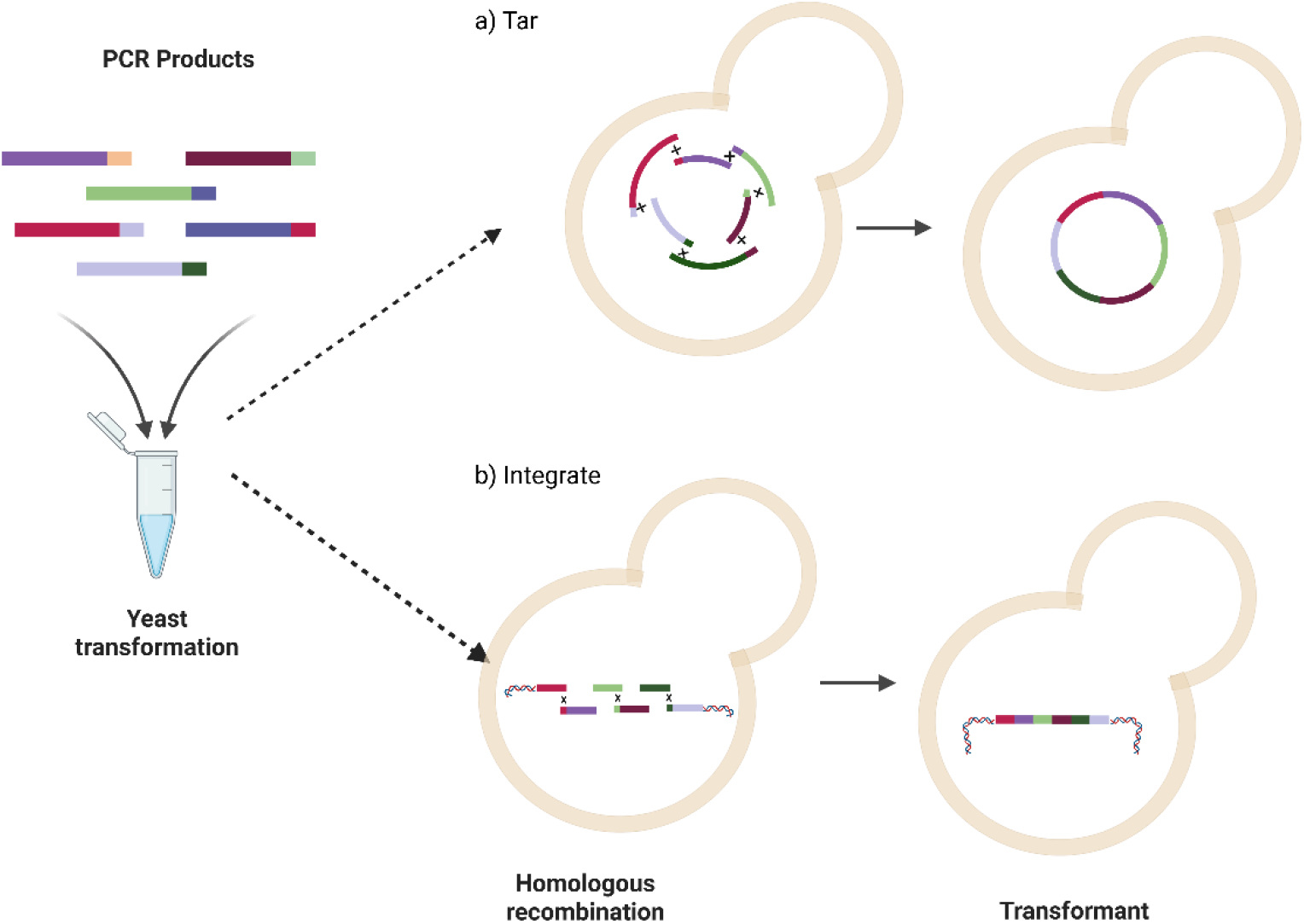
The workflow for the *tar* (a) and *integrate* (b) commands, showing initial transformation with PCR products followed by in vivo homologous recombination to assemble the final construct.

The tar command first asks the user to select a directory from among those stored in .*/data/component libraries*. These directories contain a list of components stored as .*fasta* sequences. The selection of sequences included in each library is left to the user. However, libraries may be constructed as needed for different projects, species or users by simply using the file system. The user then selects the desired components and PYEAST designs the necessary primers to amplify them, adds homology to adjacent components for assembly, and provides instructions for PCR amplification. Once this is complete, the PCR products can be directly transformed into *S. cerevisiae* without purification,^3^ where they will be assembled into a plasmid (figure 1A). Including appropriate selection markers, and origins of replication during the design stage is essential for generating plasmids suited to downstream applications. For example, if the plasmid is to be recovered and amplified in *E. coli*, the design must include a bacterial selection marker and origin of replication.

The *integrate* command functions similarly to *tar*, allowing the user to select components from the same set of libraries and design primers that introduce homology for assembly during transformation. Additionally, it includes a step where the user specifies the genomic locus of sequence insertion. These are stored as .*fasta* files, with separate upstream and downstream sequences flanking the region between which the selected sequences are assembled. Some of these include selection markers, but it is left to the user to select the appropriate combination of sequence and strains. The examples provided use integration sites derived from a widely used set of integrative plasmids.^16^ Additionally, alternate genomic sites can be added to the directory as needed.

For both *tar* and *integrate*, once components are selected and primers are designed, the .*data/primer* and .*data/templates* directories are scanned for existing primers and templates respectively. These details are then included in the instructions printed to the terminal. Upon user confirmation, PYEAST generates and saves a DNA map, an annotated .*gb* file, a complete list of primers, a list of missing primers (any primers not in the primer directory) and instructions for the necessary PCRs are saved to the output directory. The *batch* command can be used to regenerate new instruction sets for plasmids and linear assemblies stored in the output directory. Additionally, this command can create instructions for multiple assemblies. Optionally, it can also be used to generate instructions for liquid handling robots.

### Delete and Replace

The *delete* command is designed to remove arbitrary sequences from the genome using an adaptation of a method first reported by Akada et al in 2006,^5^. This approach leverages counterselection of the URA3 marker by fluoroorotic acid (FOA) and recombination between tandem repeats to remove sequences without leaving any extraneous DNA, such as antibiotic resistance genes or *loxP* sites, in the genome. The command is designed to select sequences so that, after marker removal with FOA, only the input sequence is removed. This enables precise genomic edits, such as in-frame deletions within a coding region, or partial removal of promoter elements.

The user is first prompted to input a DNA sequence to be removed, after which the command locates this sequence in the genome and designs a deletion cassette. The program also generates a set of screening primers flanking the deletion locus and provides the expected PCR product sizes for verification (figure s19). The primers, an annotated .*gb* file, a .*fasta* file, and a DNA map are then saved to the output directory.

The deletion cassette consists of: (1) upstream homology, (2) a downstream repeat to facilitate marker removal, (3) the URA3 marker, and (4) internal downstream homology within the target sequence. This cassette can be obtained as synthetic DNA, transformed directly into *S. cerevisiae*, with transformants selected on uracil drop out media. After verifying that the initial transformants contain the URA3 marker using PCR, the marker can be removed by counter selection with FOA. Recombination between the tandem repeats flanking the URA3 marker results in the complete removal of the marker and the target sequence, ensuring that only the user specified sequence is deleted (figure 2A).

**Figure 2:**
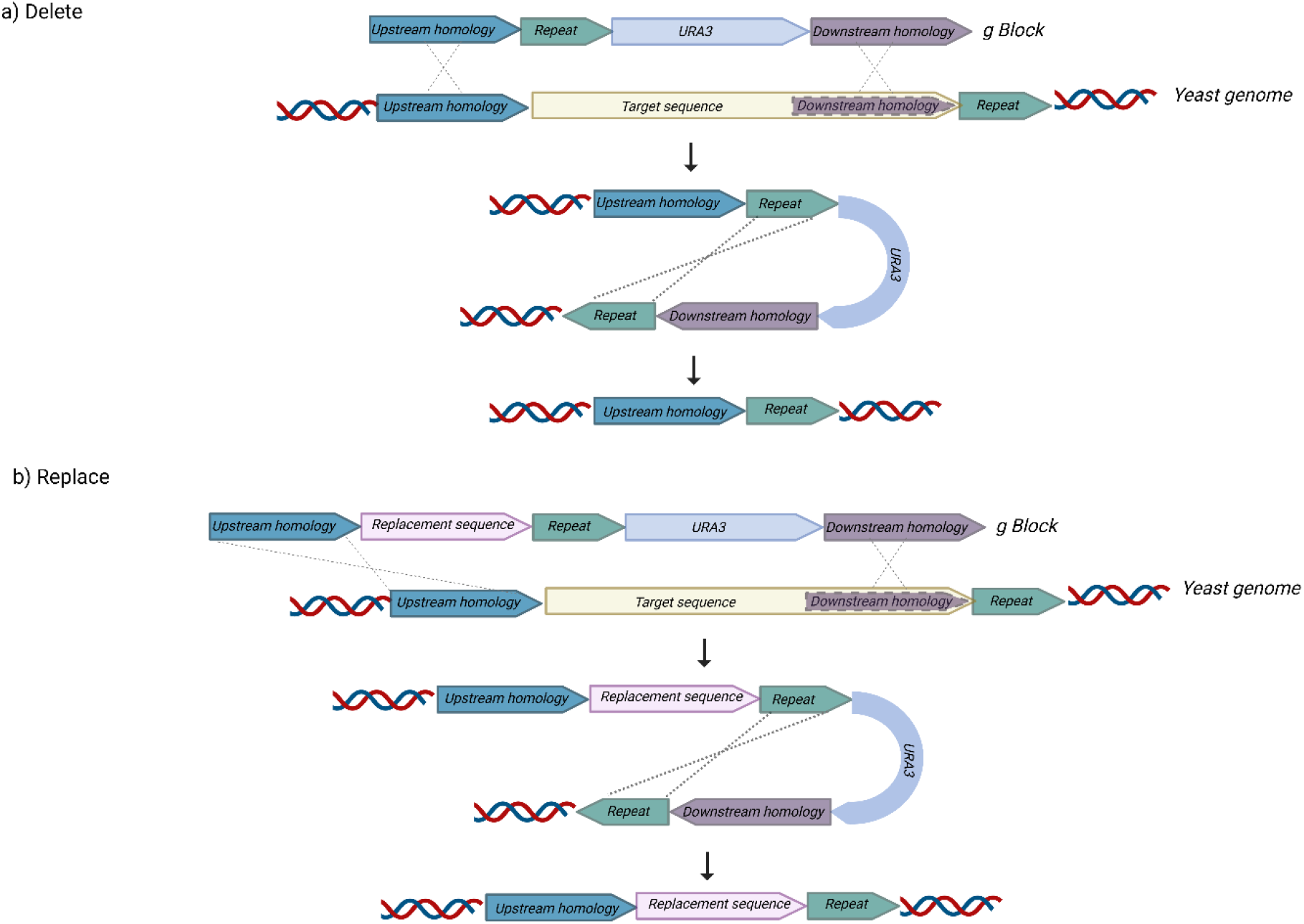
The workflow for the *delete* (a) and *replace*(b) commands, showing initial transformation with synthetic DNA fragment and URA3 marker removal by counter selection with FOA.

The *replace* command functions similarly to the *delete* command but adds a user selected sequence from one of the component folders. The user is prompted to select the position of the URA3 marker, either upstream or downstream of the replacement sequence, as this may affect the functionality of the intermediate strain prior to marker recovery. For example, when replacing the promoter of an essential gene, placing the URA3 marker upstream ensures it does not interfere with the new promoter and the coding sequence, preventing potential disruptions to gene expression that could render the intermediate strain inviable (figure 2B).

For both the *delete* and *replace* commands, the user can optionally configure the genome sequence, the file containing the URA3 marker, the repeat length, and each of the homology sequences each time the command is executed. Based on our experience the optimal lengths are highly dependent on the locus and the replacement sequence, which aligns with previous reports on how local sequence features influences the mutation rates in *S. cerevisiae*.^17,18^ We provide default parameters optimized over the course of our work,

Both commands save an annotated .*gb* file and .*fasta* file of the cassette sequence, a DNA map, and a pair of PCR screening primers to the output directory. Additionally, the expected PCR product sizes for each step of the transformation are also calculated and printed to the terminal.

## Conclusion

PYEAST addresses the gap in digital tools for *S. cerevisiae* synthetic biology between complex software packages for high throughput automated assembly strategies and more labour-intensive ad-hoc approaches common in many laboratories. By prioritizing accessibility and flexibility, PYEAST standardizes the commonly used genetic manipulations while maintaining compatibility with existing sequence repositories and laboratory resources. This approach improved efficiency and reproducibility in our own laboratory while providing a framework that can be readily adapted to diverse research settings, without requiring significant computational expertise or specialised equipment.

Looking ahead, we are actively developing additional features to enhance PYEAST’s utility. These include the integration of other digital frameworks for synthetic biology like the Synthetic Biology Open Language version 3 (SBOL3)^19^ and QUinable and Efficiently Editable Nucleotide sequence resources (QUEEN),^20^ as well as machine learning tools for sequence analysis and design optimization. The modular architecture of the codebase readily accommodates such extensions while maintaining backward compatibility with existing workflows. Modularity also encourages the adaptation of PYEAST to create tools for other model organisms where homologous recombination can be exploited in similar workflows.

The source code is available as an open-source project under the GpL2 license (https://doi.org/10.5281/zenodo.15393310). We invite the broader scientific community to adapt and extend PYEAST to address their own needs. Accessible and customisable, tools like PYEAST can help to democratize synthetic biology approaches across a wide range of laboratory settings.

## Supporting information

Supplimrntary Material for PYEAST

## Supplementary Material

Installation instructions

General methods

Worked examples:

1. Preparation of a set of plasmids using the *tar* and *batch* commands.
2. Integration of fluorescent proteins into the genome of *S. cerevisiae* using the *integrate* command.
3. Deletions of the Ade2 gene using the *delete* command.
4. Replacement of the promoter of the Flo1 gene using the *replace* command.

## Acknowledgments

A template for the Python code incorporating UV for package management was provided by Sam West (CSIRO, Energy Research Unit). We thank Oliver Mead and Bingyin Peng for initial discussions on the molecular biology of yeast; Michael Kuiper (Google DeepMind) for encouragement to publicize the codebase; Ema Johnston and Christina Gregg for a critical reading of the manuscript. This work was performed on the traditional lands of the Ngunnawal and Ngambri people.

## Author Contributions

A. M., A. S., D. V. R., M. N., and A.P. tested code and validated methods. T. L., conceived the study, wrote the code, designed and validated methods. T.L. and C. H. supervised the work. T.L. and A. S. wrote the manuscript. All authors read and approved the final manuscript.

## Notes

The authors declare no competing financial interest.

## References

(1) Tartik, M. The Priority of Yeast to Select among Various DNA Options to Repair Genome Breaks by Homologous Recombination. Mol Biol Rep 2024, 51 (1), 99. 10.1007/s11033-023-09058-0.

(2) Kuijpers, N. G.; Solis-Escalante, D.; Bosman, L.; van den Broek, M.; Pronk, J. T.; Daran, J.-M.; Daran-Lapujade, P. A Versatile, Efficient Strategy for Assembly of Multi-Fragment Expression Vectors in Saccharomyces Cerevisiae Using 60 Bp Synthetic Recombination Sequences. Microbial Cell Factories 2013, 12 (1), 47. 10.1186/1475-2859-12-47.

(3) Lin, Q.; Jia, B.; Mitchell, L. A.; Luo, J.; Yang, K.; Zeller, K. I.; Zhang, W.; Xu, Z.; Stracquadanio, G.; Bader, J. S.; Boeke, J. D.; Yuan, Y.-J. RADOM, an Efficient In Vivo Method for Assembling Designed DNA Fragments up to 10 Kb Long in Saccharomyces Cerevisiae. ACS Synth. Biol. 2015, 4 (3), 213–220. 10.1021/sb500241e.

(4) Shao, Z.; Zhao, H.; Zhao, H. DNA Assembler, an in Vivo Genetic Method for Rapid Construction of Biochemical Pathways. Nucleic Acids Research 2009, 37 (2), e16. 10.1093/nar/gkn991.

(5) Akada, R.; Kitagawa, T.; Kaneko, S.; Toyonaga, D.; Ito, S.; Kakihara, Y.; Hoshida, H.; Morimura, S.; Kondo, A.; Kida, K. PCR-Mediated Seamless Gene Deletion and Marker Recycling in Saccharomyces Cerevisiae. Yeast 2006, 23 (5), 399–405. 10.1002/yea.1365.

(6) Horecka, J.; Davis, R. W. The 50:50 Method for PCR-Based Seamless Genome Editing in Yeast. Yeast 2014, 31 (3), 103–112. 10.1002/yea.2992.

(7) Postma, E. D.; Dashko, S.; van Breemen, L.; Taylor Parkins, S. K.; van den Broek, M.; Daran, J.-M.; Daran-Lapujade, P. A Supernumerary Designer Chromosome for Modular in Vivo Pathway Assembly in Saccharomyces Cerevisiae. Nucleic Acids Research 2021, 49 (3), 1769–1783. 10.1093/nar/gkaa1167.

(8) Gibson, D. G.; Benders, G. A.; Axelrod, K. C.; Zaveri, J.; Algire, M. A.; Moodie, M.; Montague, M. G.; Venter, J. C.; Smith, H. O.; Hutchison, C. A. One-Step Assembly in Yeast of 25 Overlapping DNA Fragments to Form a Complete Synthetic Mycoplasma Genitalium Genome. Proceedings of the National Academy of Sciences 2008, 105 (51), 20404–20409. 10.1073/pnas.0811011106.

(9) Hutchison, C. A.; Chuang, R.-Y.; Noskov, V. N.; Assad-Garcia, N.; Deerinck, T. J.; Ellisman, M. H.; Gill, J.; Kannan, K.; Karas, B. J.; Ma, L.; Pelletier, J. F.; Qi, Z.-Q.; Richter, R. A.; Strychalski, E. A.; Sun, L.; Suzuki, Y.; Tsvetanova, B.; Wise, K. S.; Smith, H. O.; Glass, J. I.; Merryman, C.; Gibson, D. G.; Venter, J. C. Design and Synthesis of a Minimal Bacterial Genome. Science 2016, 351 (6280), aad6253. 10.1126/science.aad6253.

(10) Lee, M. E.; DeLoache, W. C.; Cervantes, B.; Dueber, J. E. A Highly Characterized Yeast Toolkit for Modular, Multipart Assembly. ACS Synth. Biol. 2015, 4 (9), 975–986. 10.1021/sb500366v.

(11) Panagopoulos, A.; Sideri, K. Prospect Patents and CRISPR; Rivalry and Ethical Licensing in a Semi-Commons Environment. Journal of Law and the Biosciences 2021, 8 (2), sab031. 10.1093/jlb/lsab031.

(12) Grobler, L.; Suleman, E.; Thimiri Govinda Raj, D. B. Chapter Seven - Patents and Technology Transfer in CRISPR Technology. In Progress in Molecular Biology and Translational Science; Singh, V., Ed.; Reprogramming the Genome: Applications of CRISPR-Cas in Non-mammalian Systems Part B; Academic Press, 2021; Vol. 180, pp 153–182. 10.1016/bs.pmbts.2021.01.009.

(13) Shi, Z.; Vickers, C. E. Molecular Cloning Designer Simulator (MCDS): All-in-One Molecular Cloning and Genetic Engineering Design, Simulation and Management Software for Complex Synthetic Biology and Metabolic Engineering Projects. Metabolic Engineering Communications 2016, 3, 173–186. 10.1016/j.meteno.2016.05.003.

(14) Kang, D. H.; Ko, S. C.; Heo, Y. B.; Lee, H. J.; Woo, H. M. RoboMoClo: A Robotics-Assisted Modular Cloning Framework for Multiple Gene Assembly in Biofoundry. ACS Synth. Biol. 2022, 11 (3), 1336–1348. 10.1021/acssynbio.1c00628.

(15) Nava, A. A.; Fear, A. L.; Lee, N.; Mellinger, P.; Lan, G.; McCauley, J.; Tan, S.; Kaplan, N.; Goyal, G.; Coates, R. C.; Roberts, J.; Johnson, Z.; Hu, R.; Wu, B.; Ahn, J.; Kim, W. E.; Wan, Y.; Yin, K.; Hillson, N.; Haushalter, R. W.; Keasling, J. D. Automated Platform for the Plasmid Construction Process. ACS Synth. Biol. 2023, 12 (12), 3506–3513. 10.1021/acssynbio.3c00292.

(16) Gnügge, R.; Liphardt, T.; Rudolf, F. A Shuttle Vector Series for Precise Genetic Engineering of Saccharomyces Cerevisiae. Yeast 2016, 33 (3), 83–98. 10.1002/yea.3144.

(17) Zhu, Y. O.; Siegal, M. L.; Hall, D. W.; Petrov, D. A. Precise Estimates of Mutation Rate and Spectrum in Yeast. Proceedings of the National Academy of Sciences 2014, 111 (22), E2310–E2318. 10.1073/pnas.1323011111.

(18) Kiktev, D. A.; Sheng, Z.; Lobachev, K. S.; Petes, T. D. GC Content Elevates Mutation and Recombination Rates in the Yeast Saccharomyces Cerevisiae. Proceedings of the National Academy of Sciences 2018, 115 (30), E7109–E7118. 10.1073/pnas.1807334115.

(19) Mitchell, T.; Beal, J.; Bartley, B. pySBOL3: SBOL3 for Python Programmers. ACS Synth. Biol. 2022, 11 (7), 2523–2526. 10.1021/acssynbio.2c00249.

(20) Mori, H.; Yachie, N. A Framework to Efficiently Describe and Share Reproducible DNA Materials and Construction Protocols. Nat Commun 2022, 13 (1), 2894. 10.1038/s41467-022-30588-x.

