## Supplementary material for "PYEAST – Python Enabled Automated Strain Transformation": Supplimrntary Material for PYEAST

1. CSIRO, Agriculture and Food, Canberra, 2601, Australia.
2. Department of Microbiology, Faculty of Life Sciences, Ahmadu Bello University, Zaria 810107, Nigeria

**Installation instructions**

**Prerequisites**

uv, git

To install uv, see the instructions: https://docs.astral.sh/uv/getting-started/installation/, in short:

In bash on (most) Linux systems and Mac:

```
curl -LsSf https://astral.sh/uv/0.4.6/install.sh | sh
```

Restart your shell and make sure ` uv --version` works

In Windows, from any shell run:

```
powershell -ExecutionPolicy ByPass -c "irm https://astral.sh/uv/install.ps1 | iex"
```

Restart your shell and make sure ` uv --version` works

Git can be installed by following the instructions available at:

https://github.com/git-guides/install-git

To install PYEAST, first Clone the repository

In bash on (most) Linux systems, Mac terminal or windows cmd or git bash run:

```
git clone https://github.com/TomLoan/PYEAST
```

In the same shell, navigate into the director where PYEAST was installed by default this is PYEAST:

```
cd pyeast
```

then, still in the same shell run PYEAST using uv:

```
uv run pyeast
```

uv will handle the package management and create the required virtual environment in the local directory.

Once this process is complete a range of commands will be printed to the terminal, use uv run pyeast command --help for more information on running each command

**General Methods.**

Chemical reagents were purchased from Merck (NJ, USA), DNA polymerase was purchased from New England Biolabs (Ma, USA). *Saccharomyces cerevisiae* strain BY4741 (Mata his3Δ1 leu2Δ0 met15Δ0 ura3Δ0) (ATCC no. 201388) was used for all examples described below. BY4741 was propagated on YPD media (1% w/v Bacto yeast extract, 2% w/v Bacto peptone, 2% w/v glucose) or synthetic drop-out media (SD) without either leucine, histidine or uracil (0.67% w/v yeast nitrogen base (YNB) without amino acids, 0.162% w/v yeast synthetic dropout media supplement without leucine (Merck Y1376), or 0.192% w/v supplement without histidine (Merck Y1751) or 0.192% w/v supplement without uracil (Merck Y1501), 2% w/v glucose) as required. Yeast was transformed using the yeast transformation kit (Merck yeast1) according to the manufacturer's instructions.

**Worked examples**

**Assembly of a set of plasmids using the tar and batch commands.**

1. In shell program (powershell, cmd on windows, bash on linux or terminal on Mac), navigate to the directory where the git repository was cloned into and run the tar command in a terminal (figure S1).

```
PS C:\Users\loa012\Python\PYEAST> uv run pyeast tar
```

Figure S1. Running the tar command in a terminal using uv.

2. Select the Saccharomyces cerevisiae component library. Use tab to auto complete the selection (figure 2A). Use the enter key to confirm selection. This prints a table of available sequences to the terminal (figure S2B).

a)

```
data\component libraries
    Available Component
        Directories

 Name

 Saccharomyces cerevisiae

Select components directory (TAB for suggestions): Saccharomyces cerevisiae
                                                     Saccharomyces cerevisiae
```

b)

| Name | Length | Description |
|---|---|---|
| 2Micron | 1343 bp | 2Micron S.cerevisiae high copy replication origin. Seqeunce taken from pESC plasmid |
| 2Micron_pYES2 | 881 bp | 2Micron_pYES2 2micron origin from pYes2 vector shortwe than pESC |
| ampR | 1086 bp | ampR ampicillian resistance cassette from pUC19 |
| AmpR_ColE1 | 1769 bp | AmpR_ColE1 Ampicillin resistance gene and ColE1 origin of replication (high copy) |
| CEN6_ARS4 | 519 bp | CEN6_ARS4 Origin of replication, low copy |
| CmR_p15A | 1955 bp | CmR_p15A Chloramphenicol resistance gene and p15A origin of replication |
| ColE1 | 683 bp | ColE1 replication origin from pUC19 |
| ERG10 | 1197 bp | ERG10 YPL028W SGDID:S000005949, Chromosome XVI:498096..499292 |
| His3 | 1106 bp | His3 from pUC19-Pot1::His3 -Tom L |
| hphMX6 | 1576 bp | hphMX6 Cassette confers hygromycin resistance in S. cerevisiae |
| kanMX | 1357 bp | kanMX confers resistance to genticin (G418) in S. cerevisiae |
| KanR_ColE1 | 1649 bp | KanR_ColE1 Kanamycin resistance gene and ColE1 origin of replication |
| Leu2d | 1194 bp | Leu2d Leu2 marker with truncated promoter to decrease expression for increased plasmid copy number or integration events |
| Leu2Mx | 2265 bp | Leu2Mx gene from K. lactis, complements leu2 allele in S. cerevisiae |
| mCherry | 711 bp | mCherry Red Fluorescent protein, codon optimised for S. cerevisiae |
| moxBFP | 720 bp | moxBFP Blue fluorescent protein codon optimised for S. cerevisiae |
| PBR322 | 620 bp | PBR322 bacterial replication origin |
| pCTT1 | 1000 bp | pCTT1 Stress responsive promoter from S. cerevisiae gene: YGR088W |
| pDDR2 | 1000 bp | pDDR2 Stress repsonsive promoter from S. cerevisiae gene: YOL052C-A |
| PDI1 | 1569 bp | PDI1 protein disulfide isomerase from S. cerevisiae gene: YCL043C SGDID:S000000548, Chromosome III:48653..50221 |
| pDOG1 | 1000 bp | pDOG1 promter regulated by HAC1 from Sc288 gDNA |
| pFMP16 | 1000 bp | pFMP16 Stress repsonsive promoter from S. cerevisiae gene: YDR070C |
| pGal1 | 442 bp | pGal1 Galactose responsive promoter. Sequence taken from the pYES2 vector |
| pGal1_10 | 665 bp | pGal1_10 bidirectional Gal promoter. Sequence taken from pESC plasmid |

Figure S2. A) Selection of the components directory. Note additional folders containing components as *.fasta* files will be displayed in the list of available component libraries. These can be added and modified using the file browser. B) First 24 lines of the list of components inside the Saccharomyces cerevisiae component library.

3. Select components in the order required, separate each selection with “space” (figure S3). The “tab” key can be used to select options for autocompletion. A numbered list of the selections made will be printed to the terminal, selection can be confirmed by pressing ‘y’.

```
Enter sequences to assemble (space-separated)
Use TAB for autocompletion
Sequences: pDOG1 SHU1 tDIT1 pMTR10 SHU2 tLSC2 pKEG1 CSM2 tRPL3 pVPS17 PSY3 tIDP1 Leu2Mx AmpR_ColE1 2Micron_pYES2
                                                                                                    2Micron
                                                                                                    2Micron_pYES2
```

Figure S3 component selection. In this example several genes from the homologous recombination pathway in S. cerevisiae are selected for expression under weak promoters.

4. PCR instructions will be printed to the terminal (figure S4), including primers present in *.data/primers* and templates identified from *.data/templates* and the expected size of each PCR product in base pairs. Primers shown here are identified by sequence from *.xlsx* files stored in *./data/primers* (See note 3), and templates are identified from sequence files (*.gb* or *.fasta*) stored in *.data/templates*.

```
Designing primers...
Checking primer locations...
Finding templates...
Rationalizing selections...
Generating instructions...
```

*Assembly Instructions*

| Part Name | Fwd Primer | Fwd Plate | Well | Rev Primer | Rev Plate | Well | Template | Size |
|---|---|---|---|---|---|---|---|---|
| pDOG1 | pDOG1_0F-2Micron_pYES2 | Demo Plate | A1 | pDOG1_0R-SHU1 | Demo Plate | A2 | chr08 | 1050 |
| SHU1 | SHU1_1F-pDOG1 | Demo Plate | A3 | SHU1_1R-tDIT1 | Demo Plate | A4 | chr08 | 503 |
| tDIT1 | tDIT1_2F-SHU1 | Demo Plate | A5 | tDIT1_2R-pMTR10 | Demo Plate | A6 | chr04 | 484 |
| pMTR10 | pMTR10_3F-tDIT1 | Demo Plate | A7 | pMTR10_3R-SHU2 | Demo Plate | A8 | chr15 | 1050 |
| SHU2 | SHU2_4F-pMTR10 | Demo Plate | A9 | SHU2_4R-tLSC2 | Demo Plate | A10 | chr04 | 722 |
| tLSC2 | tLSC2_5F-SHU2 | Demo Plate | A11 | tLSC2_5R-pKEG1 | Demo Plate | A12 | chr07 | 550 |
| pKEG1 | pKEG1_6F-tLSC2 | Demo Plate | B1 | pKEG1_6R-CSM2 | Demo Plate | B2 | chr06 | 1050 |
| CSM2 | CSM2_7F-pKEG1 | Demo Plate | B3 | CSM2_7R-tRPL3 | Demo Plate | B4 | chr09 | 692 |
| tRPL3 | tRPL3_8F-CSM2 | Demo Plate | B5 | tRPL3_8R-pVPS17 | Demo Plate | B6 | chr15 | 543 |
| pVPS17 | pVPS17_9F-tRPL3 | Demo Plate | B7 | pVPS17_9R-PSY3 | Demo Plate | B8 | chr15 | 990 |
| PSY3 | PSY3_10F-pVPS17 | Demo Plate | B9 | PSY3_10R-tIDP1 | Demo Plate | B10 | chr12 | 779 |
| tIDP1 | tIDP1_11F-PSY3 | Demo Plate | B11 | tIDP1_11R-Leu2Mx | Demo Plate | B12 | chr04 | 512 |
| Leu2Mx | Leu2Mx_12F-tIDP1 | Demo Plate | C1 | Leu2Mx_12R-AmpR_ColE1 | Demo Plate | C2 | pTw_Leu2MX | 2315 |
| AmpR_ColE1 | AmpR_ColE1_13F-Leu2Mx | Demo Plate | C3 | AmpR_ColE1_13R-2Micron_pYES2 | Demo Plate | C4 | pUC19 | 1819 |
| 2Micron_pYES2 | 2Micron_pYES2_14F-AmpR_ColE1 | Demo Plate | C5 | 2Micron_pYES2_14R-pDOG1 | Demo Plate | C6 | pYES2 | 931 |

```
Proceed with assembly? [y/N]:
```

Figure S4. PCR instructions for the selected design. Note primers and templates are selected by sequence matching from sequences stored in *.data/primers* and *.data/templates* respectively.

5. Input a name for the plasmid, this name will be used for the sequence file (*.gb* locus) and the file names (figure S5).

```
Enter a name prefix for your output files
Files will be saved in the output directory
Name prefix: pLeu2_Shu_2u
output\pLeu2_Shu_2u
Generating assembly...
Saving outputs...
Files saved:
GenBank file: output\pLeu2_Shu_2u.gb
Sequence map: output\pLeu2_Shu_2u_map.png
Assembly instructions: output\pLeu2_Shu_2u_instructions.tsv
All primers: output\pLeu2_Shu_2u_all_primers.tsv
Saving outputs...
✓ Plasmid design complete!
Saving outputs...
```

Figure S5. Outputs are saved to *.data/output*.

6. Design other plasmids as required, in this example a series of four related plasmids is designed, each containing either 15 (pLeu2_Shu_2u and pLeu2_Shu_CEN), or 18 different components (pLeu2_Shu_RAD51_2u and pLeu2_SHU_RAD51).

7. Once all plasmid designs are complete, primers and templates should be acquired and added to the appropriate folders in *./data*. To re-generate the instructions for all the plasmids run the batch command from the directory where PYEAST is stored by inputting "uv run pyeast" in a terminal and select the vectors to be assembled (figure S6). The "tab" key can be used for autocompletion. In this example we select the four related plasmids designed in the above steps. Note that the high degree of similarity allows the reuse of many of the PCR products and primers, so relatively few additional reactions and primers are required to assemble the additional vectors. Note that the batch command can accept both circular constructs designed using the tar command and linear constructs designed using the integrate command.

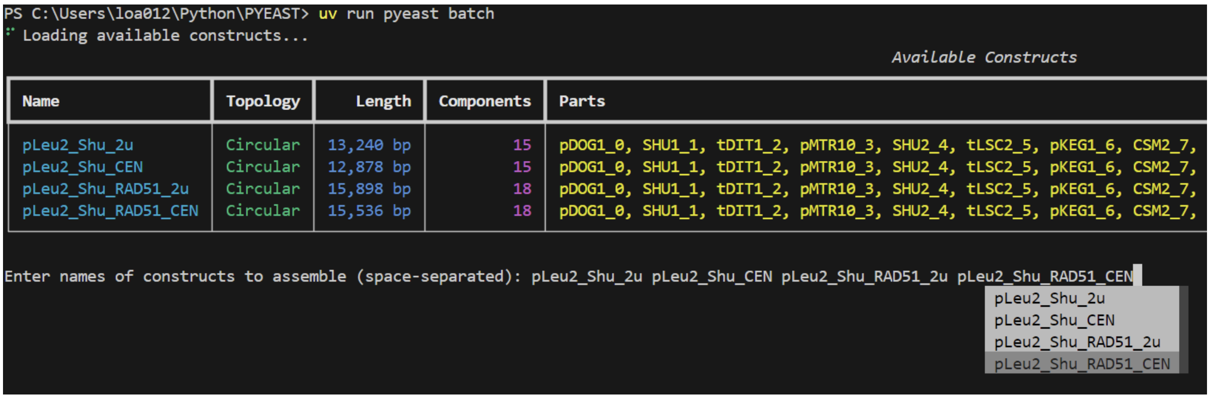


Figure S6. Running the batch command using uv and select the constructs to be assembled in this batch.

8. Information about the necessary PCRs, including the number of unique reactions, primers and templates identified will be printed to the terminal. Confirm with "y+enter". Input a name for the batch (figure S7).

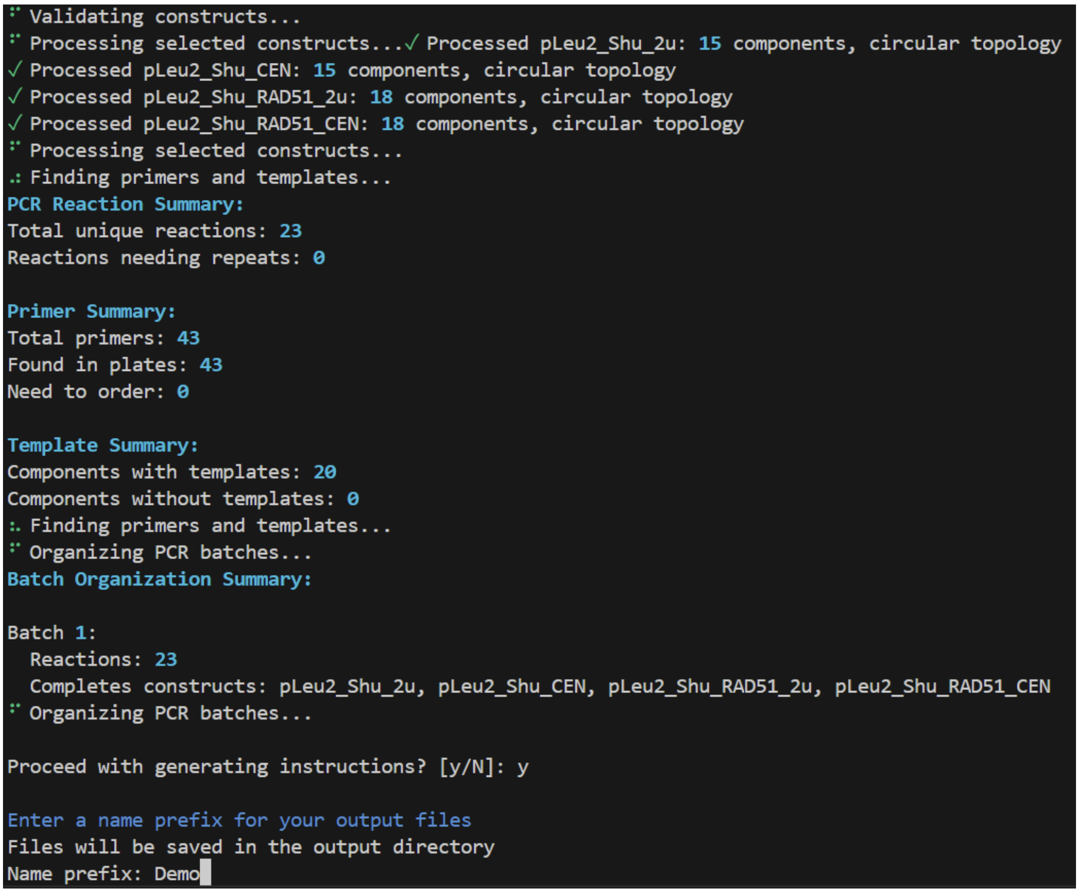


Figure S7. Preliminary information on the selected batch printed to the terminal.

9. Instructions for the necessary PCRs and which reactions are required for each construction will be printed to the terminal (figure S8) and saved to *./output*. Note, in this example many of the PCR products are required for some or all of all the assemblies, but as only 2 µL of each reaction is required for transformation they are only run once. There is an option for the batch command (--re_use_limit) that controls how many times a PCR can be reused in assemblies before it is run again. By default, the re_use_limit is set to 5. For details on how to use this option input uv run pyeast batch --help in a terminal.

Assembly Instructions:

Assembly Instructions

| Batch | Part Name | Constructs | Fwd Primer | Fwd Plate | Well | Rev Primer | Rev Plate | Well | Template | Size | Repeat |
|---|---|---|---|---|---|---|---|---|---|---|---|
| 1 | pDOG1_0 | pLeu2_Shu_2u, pLeu2_Shu_RAD51_2u | pDOG1_0F-2Micron_pYES2_forward | Demo Plate | A1 | pDOG1_0R-SHU1_reverse | Demo Plate | A2 | chr08 | 1074 | No |
| 1 | SHU1_1 | pLeu2_Shu_2u, pLeu2_Shu_CEN, pLeu2_Shu_RAD51_2u, pLeu2_Shu_RAD51_CEN | SHU1_1F-pDOG1_forward | Demo Plate | A3 | SHU1_1R-tDIT1_reverse | Demo Plate | A4 | chr08 | 503 | No |
| 1 | tDIT1_2 | pLeu2_Shu_2u, pLeu2_Shu_CEN, pLeu2_Shu_RAD51_2u, pLeu2_Shu_RAD51_CEN | tDIT1_2F-SHU1_forward | Demo Plate | A5 | tDIT1_2R-pMTR10_reverse | Demo Plate | A6 | chr04 | 503 | No |
| 1 | pMTR10_3 | pLeu2_Shu_2u, pLeu2_Shu_CEN, pLeu2_Shu_RAD51_2u, pLeu2_Shu_RAD51_CEN | pMTR10_3F-tDIT1_forward | Demo Plate | A7 | pMTR10_3R-SHU2_reverse | Demo Plate | A8 | chr15 | 1050 | No |
| 1 | SHU2_4 | pLeu2_Shu_2u, pLeu2_Shu_CEN, pLeu2_Shu_RAD51_2u, pLeu2_Shu_RAD51_CEN | SHU2_4F-pMTR10_forward | Demo Plate | A9 | SHU2_4R-tLSC2_reverse | Demo Plate | A10 | chr04 | 722 | No |
| 1 | tLSC2_5 | pLeu2_Shu_2u, pLeu2_Shu_CEN, pLeu2_Shu_RAD51_2u, pLeu2_Shu_RAD51_CEN | tLSC2_5F-SHU2_forward | Demo Plate | A11 | tLSC2_5R-pKEG1_reverse | Demo Plate | A12 | chr07 | 550 | No |
| 1 | pKEG1_6 | pLeu2_Shu_2u, pLeu2_Shu_CEN, pLeu2_Shu_RAD51_2u, pLeu2_Shu_RAD51_CEN | pKEG1_6F-tLSC2_forward | Demo Plate | B1 | pKEG1_6R-CSM2_reverse | Demo Plate | B2 | chr06 | 1077 | No |
| 1 | CSM2_7 | pLeu2_Shu_2u, pLeu2_Shu_CEN, pLeu2_Shu_RAD51_2u, pLeu2_Shu_RAD51_CEN | CSM2_7F-pKEG1_forward | Demo Plate | B3 | CSM2_7R-tRPL3_reverse | Demo Plate | B4 | chr09 | 721 | No |
| 1 | tRPL3_8 | pLeu2_Shu_2u, pLeu2_Shu_CEN, pLeu2_Shu_RAD51_2u, pLeu2_Shu_RAD51_CEN | tRPL3_8F-CSM2_forward | Demo Plate | B5 | tRPL3_8R-pVPS17_reverse | Demo Plate | B6 | chr15 | 543 | No |
| 1 | pVPS17_9 | pLeu2_Shu_2u, pLeu2_Shu_CEN, pLeu2_Shu_RAD51_2u, pLeu2_Shu_RAD51_CEN | pVPS17_9F-tRPL3_forward | Demo Plate | B7 | pVPS17_9R-PSY3_reverse | Demo Plate | B8 | chr15 | 990 | No |
| 1 | PSY3_10 | pLeu2_Shu_2u, pLeu2_Shu_CEN, pLeu2_Shu_RAD51_2u, pLeu2_Shu_RAD51_CEN | PSY3_10F-pVPS17_forward | Demo Plate | B9 | PSY3_10R-tIDP1_reverse | Demo Plate | B10 | chr12 | 799 | No |
| 1 | tIDP1_11 | pLeu2_Shu_2u, pLeu2_Shu_CEN | tIDP1_11F-PSY3_forward | Demo Plate | B11 | tIDP1_11R-Leu2Mx_reverse | Demo Plate | B12 | chr04 | 528 | No |
| 1 | Leu2Mx_12 | pLeu2_Shu_2u, pLeu2_Shu_CEN | Leu2Mx_12F-tIDP1_forward | Demo Plate | C1 | Leu2Mx_12R-AmpR_ColE1_reverse | Demo Plate | C2 | pTw_Leu2MX | 2315 | No |
| 1 | AmpR_ColE1_13 | pLeu2_Shu_2u, pLeu2_Shu_RAD51_2u | AmpR_ColE1_13F-Leu2Mx_forward | Demo Plate | C3 | AmpR_ColE1_13R-2Micron_pYES2_reverse | Demo Plate | C4 | pUC19 | 1819 | No |
| 1 | 2Micron_pYES2_14 | pLeu2_Shu_2u, pLeu2_Shu_RAD51_2u | 2Micron_pYES2_14F-AmpR_ColE1_forward | Demo Plate | C5 | 2Micron_pYES2_14R-pDOG1_reverse | Demo Plate | C6 | pYES2 | 949 | No |
| 1 | pDOG1_0 | pLeu2_Shu_CEN, pLeu2_Shu_RAD51_CEN | pDOG1_0F-CEN6_ARS4_forward | Demo Plate | C7 | pDOG1_0R-SHU1_reverse | Demo Plate | A2 | chr08 | 1074 | No |
| 1 | AmpR_ColE1_13 | pLeu2_Shu_CEN, pLeu2_Shu_RAD51_CEN | AmpR_ColE1_13F-Leu2Mx_forward | Demo Plate | C3 | AmpR_ColE1_13R-CEN6_ARS4_reverse | Demo Plate | C8 | pUC19 | 1819 | No |
| 1 | CEN6_ARS4_14 | pLeu2_Shu_CEN, pLeu2_Shu_RAD51_CEN | CEN6_ARS4_14F-AmpR_ColE1_forward | Demo Plate | C9 | CEN6_ARS4_14R-pDOG1_reverse | Demo Plate | C10 | pRS316 | 569 | No |
| 1 | tIDP1_11 | pLeu2_Shu_RAD51_2u, pLeu2_Shu_RAD51_CEN | tIDP1_11F-PSY3_forward | Demo Plate | B11 | tIDP1_11R-pSIL1_reverse | Demo Plate | C11 | chr04 | 528 | No |
| 1 | pSIL1_12 | pLeu2_Shu_RAD51_2u, pLeu2_Shu_RAD51_CEN | pSIL1_12F-tIDP1_forward | Demo Plate | C12 | pSIL1_12R-RAD51_reverse | Demo Plate | D1 | chr15 | 1072 | No |
| 1 | RAD51_13 | pLeu2_Shu_RAD51_2u, pLeu2_Shu_RAD51_CEN | RAD51_13F-pSIL1_forward | Demo Plate | D2 | RAD51_13R-tRPL41B_reverse | Demo Plate | D3 | chr05 | 1253 | No |
| 1 | tRPL41B_14 | pLeu2_Shu_RAD51_2u, pLeu2_Shu_RAD51_CEN | tRPL41B_14F-RAD51_forward | Demo Plate | D4 | tRPL41B_14R-Leu2Mx_reverse | Demo Plate | D5 | chr04 | 505 | No |
| 1 | Leu2Mx_15 | pLeu2_Shu_RAD51_2u, pLeu2_Shu_RAD51_CEN | Leu2Mx_15F-tRPL41B_forward | Demo Plate | D6 | Leu2Mx_15R-AmpR_ColE1_reverse | Demo Plate | C2 | pTw_Leu2MX | 2315 | No |

Assembly Groups:

Assembly Groups

| Batch | Construct | Required_Wells | Topology |
|---|---|---|---|
| 1 | pLeu2_Shu_2u | Well 1, Well 2, Well 3, Well 4, Well 5, Well 6, Well 7, Well 8, Well 9, Well 10, Well 11, Well 12, Well 13, Well 14, Well 15 | circular |
| 1 | pLeu2_Shu_CEN | Well 16, Well 2, Well 3, Well 4, Well 5, Well 6, Well 7, Well 8, Well 9, Well 10, Well 11, Well 12, Well 13, Well 17, Well 18 | circular |
| 1 | pLeu2_Shu_RAD51_2u | Well 1, Well 2, Well 3, Well 4, Well 5, Well 6, Well 7, Well 8, Well 9, Well 10, Well 11, Well 19, Well 20, Well 21, Well 22, Well 23, Well 14, Well 15 | circular |
| 1 | pLeu2_Shu_RAD51_CEN | Well 16, Well 2, Well 3, Well 4, Well 5, Well 6, Well 7, Well 8, Well 9, Well 10, Well 11, Well 19, Well 20, Well 21, Well 22, Well 23, Well 17, Well 18 | circular |

Figure S8. Instructions for the required PCRs of a batch, and which reactions are required for the assembly of each construct.

10. [Optional] Generate instructions for a liquid handler. The requires primers to be stored in plates, and templates to be stored and input into the *./data/tamplates/TemPlates.xlsx* file. (See note 4). On release this command only provides instructions for the epMotion 5075. Outputs for other instruments are planned and can be configured by the user if necessary. This generates a csv file for the required transfers for primers and templates for PCR set up, and a second instruction set for the combinations of PCR products required for each construct.
11. Carry out PCR according to instructions provided. A high-fidelity polymerase is recommended to avoid introducing sequence variations. We have had success with both Phusion and Q5 both from New England Biolabs (MA, USA). Amplification can be verified by gel electrophoresis as below or similar. In this example, PCRs were set up using an epMotion 5075.
12. Transform all the PCR products into *S. cerevisiae* using the PEG, LiOAc method.[1] We typically use 2 µL of each PCR product.
13. Plate cells on selective media and grow for 2-3 days at 30°C. In this example, each transformation was pelleted by centrifugation (1000 g, 1 minute), the supernatant was removed, and cells were resuspended in 100 µL of sterile water before plating on SC-Leu +2% Glucose. Plates were incubated for 3 days at 30°C, resulting in a lawn of densely packed colonies. (See Note 1)
14. Colonies from approximately 4 $cm^2$ from each transformation plate were collected and pooled for plasmid extraction using the Zymoprep yeast plasmid miniprep II kit (Zymogen, Ca, USA) according to the manufacturer's instructions. The resulting 10 µL of plasmid DNA was transformed into *E. coli* (NEB10β, NEB Ma, USA) according to the manufacturer's instructions and plated on LB supplemented with 50 mg/L ampicillin (See note 2).
15. Four *E. coli* colonies were selected at random from each plate and cultured in 5 mL of LB + 50 mg/L ampicillin for plasmid extraction using the Monarch plasmid miniprep kit (NEB) for whole plasmid sequencing.
16. Whole plasmid sequencing was performed by Plasmidsaurus using Oxford Nanopore Technology with custom analysis and annotation. For the two plasmids contain 15 distinct components (pLeu2_Shu_2u and pLeu2_Shu_CEN) all four plasmids had the correct sequences, whereas for the two plasmids with 18 distinct components 2 of the 4 plasmids analysed had the correct sequences.

**Integration of fluorescent proteins into the genome of *S. cerevisiae* using the integrate command.**

1. In shell program (powershell, cmd on windows, bash on linux or terminal on Mac), navigate to the directory where the git repository was cloned and run the integrate command in a terminal (figure S9).

```
PS C:\Users\loa012\Python\PYEAST> uv run pyeast integrate
```

Figure S9. Running the integrate command in a terminal using uv.

2. Select the Saccharomyces cerevisiae component library. Use tab to auto complete the selection. Use the enter key to confirm selection. This prints a table of available sequences (figure S2B) and integration sites (figure S10) to the terminal.

*Available Integration Sites:*

| Name | Up | Down | Description |
|---|---|---|---|
| His3MXint | 1313 bp | 307 bp | His3MX_up His3 marker from S. pombe |
| Leu2int | 1782 bp | 175 bp | Leu2int_up |
| Leu2MXint | 2478 bp | 332 bp | Leu2Mx_up Leu2 gene from K. lactis |
| Leu2Ty1d | 1582 bp | 360 bp | Leu2Ty1d_up for high copy number intergration at the Ty1delta transposon locus │ Owner Tom L |
| PDC6 | 543 bp | 469 bp | PDC6_up |
| Ura3MX | 1838 bp | 226 bp | Ura3MX_Up Ura3 gene from C. albicans |

Figure S10. Available integration sites are printed to the terminal. Not each file contains both upstream and downstream sequences, which flank the integration site and contain homology to chromosomal DNA for targeted integration. Not all integration sites include selection markers in which case they will need to be included in selected components in the previous step

3. Select components in the order required, separate each selection with “space” (figure S11). The “tab” key can be used to select options for autocompletion. In this example yeast enhanced green or red (instructions not shown) fluorescent protein (YeGFP and YeRFP) with the Tef1 promoter and RPL41A terminator are selected for insertion into the His3MX site.

```
Enter the names of the components you want to assemble, in order (space-separated): pTEF1 YeGFP tRPL41A
                                                                                                 tRPL41A
                                                                                                 tRPL15A
                                                                                                 tRPL3
                                                                                                 tRPL41B
```

Figure S11. Component selection for integration.

4. Select the integration site, as with component selection “tab” can be used to autocomplete the selection. A numbered list of the selections made will be printed to the terminal, selection can be confirmed by pressing ‘y’ (figure S12A). PCR instructions will be printed to the terminal (figure S12B), including primers present in *.data/primers* and templates identified from *.data/templates* and the expected size of each PCR product in base pairs. Primers shown here are identified by sequence from *.xlsx* files stored in *./data/primers* (See note 3), and templates are identified from sequence files (*.gb* or *.fasta*) stored in *.data/templates*.

a)

```
Enter the name of the integration site: His3MXint
Found integration site: His3MXint

Selected assembly:
Components:
1. pTEF1
2. YeGFP
3. tRPL41A
Integration site: His3MXint
Is this correct? (y/n)
```

b)

*Assembly Instructions*

| Part Name | Fwd Primer | Fwd Plate | Well | Rev Primer | Rev Plate | Well | Template | Size |
|---|---|---|---|---|---|---|---|---|
| His3MX_up | His3MX_up_F | Demo Plate | D7 | His3MX_up_R-pTEF1 | Demo Plate | D8 | pTw_His3MX | 1338 |
| pTEF1 | pTEF1_F-His3MX_up | Demo Plate | D9 | pTEF1_R-YeGFP | Demo Plate | D10 | chr16 | 462 |
| YeGFP | YeGFP_F-pTEF1 | Demo Plate | D11 | YeGFP_R-tRPL41A | Demo Plate | D12 | YeGFP-pTwistAmp-Highcopy | 770 |
| tRPL41A | tRPL41A_F-YeGFP | Demo Plate | E1 | tRPL41A_R-His3MX_down | Demo Plate | E2 | chr04 | 525 |
| His3MX_down | His3MX_down_F-tRPL41A | Demo Plate | E3 | His3MX_down_R | Demo Plate | E4 | pTw_His3MX | 332 |

Figure S12. A) confirmation of components and integration site selected for assembly. B) PCR instructions for YeGFP integration.

5. Carry out PCR according to instructions provided. A high-fidelity polymerase is recommended to avoid introducing sequence variations. We have had success with both Phusion and Q5 both from New England Biolabs (MA, USA). Amplification can be verified by gel electrophoresis as below or similar.
6. Transform all the PCR products into *S. cerevisiae* using the PEG, LiOAc method.[1] We typically use 2 µL of each PCR product.
7. Plate cells on selective media and grow for 2-3 days at 30°C. In this example, 10 % of the transformation was plated directly on SC-His + 2% Glucose. The remaining cells from each transformation were pelleted by centrifugation (1000 g, 1 minute), the supernatant was removed, and cells were resuspended in 100 µL of sterile water before plating on SC-His +2% Glucose. Plates were incubated for 3 days at 30°C.
8. Five randomly selected colonies were screened by PCR using His3MX_up_F and His3MX_down_R primers. All colonies screened contained the correctly sized PCR product (3.2 kb) as compared to the wild type (800 bp). Expression of fluorescent proteins was confirmed by fluorescent microscopy (figure S13B).

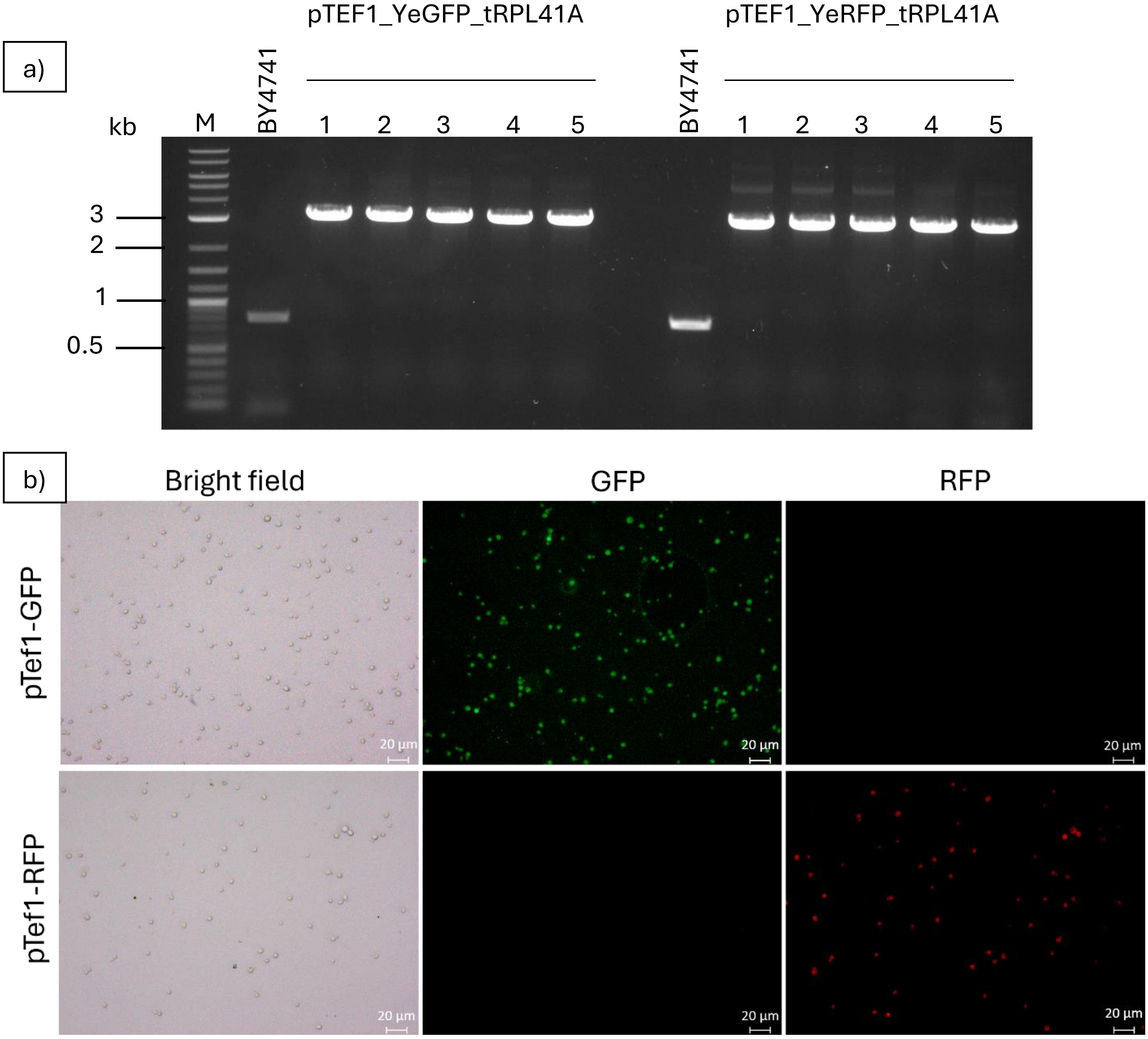


Figure S13A) Gel electrophoresis confirming integration of the correct size. Marker (Quick-Load® Purple 1 kb Plus DNA Ladder, New England Biolabs (MA, USA)); BY4741 (parent strain); pTEF1_YeGFP_tRPL41A colonies 1-5; pTef1_YeRFP_tRPL41A colonies 1-5. B) Fluorescent microscope images of cells containing YeGFP (top) and cells containing YeRFP (bottom).

**Deletion of the Ade2 gene using the delete command**

1. In shell program (powershell, cmd on windows, bash on linux or terminal on Mac), navigate to the directory where the git repository was cloned and run the delete command (figure S14). In this example we use the command options to change the repeat length to 40 bp and the downstream homology length to 60 bp.

```
PS C:\Users\loa012\Python\PYEAST> uv run pyeast delete --downstream_homology_len 60 --repeat_length 40
```

Figure S14. Running the delete command in a terminal using uv. Note the downstream_homology_length and repeat_length options are used to change the size of these features from their default values (200 and 160 bp respectively)

2. Input the deletion sequence (figure S15). Note care must be taken not to copy in return keys, which are present in *.fasta* files. Typically, we copy from a sequence browser or similar application (geneious, benchling, snapgene etc). In this example we input the coding region for the ADE2 gene. The input sequence should be long enough to be unique and should be longer than the downstream homology length. The sequence should exactly match part of the genome. For strains other than the default (BY4741) the genome_file option can be used to input the path to the corresponding genome file.

```
Enter the DNA sequence you want to delete:
Use only A, T, G, and C
Sequence: ATGGATTCTAGAACAGTTGGTATATTAGGAGGGGACAATTGGGACGTATGATTGTTGAGGCAGCAAACAGGCTCAACATTAAGA
CGGTAATACTAGATGCTGAAAATTCTCCTGCCAAACAAATAAGCAACTCCAATGACCACGTTAATGGCTCCTTTTCCAATCCTCTTGATATCGAA
AAACTAGCTGAAAAATGTGATGTGCTAACGATTGAGATTGAGCATGTTGATGTTCCTACACTAAAGAATCTTCAAGTAAAACATCCCAAATTAAA
AATTTACCCTTCTCCAGAAACAATCAGATTGATACAAGACAAATATATTCAAAAAGAGCATTTAATCAAAAATGGTATAGCAGTTACCCAAAGTG
TTCCTGTGGAACAAGCCAGTGAGACGTCCCTATTGAATGTTGGAAGAGATTTGGGTTTTCCATTCGTCTTGAAGTCGAGGACTTTGGCATACGAT
GGAAGAGGTAACTTCGTTGTAAAGAATAAGGAAATGATTCCGGAAGCTTTGGAAGTACTGAAGGATCGTCCTTTGTACGCCGAAAAATGGGCACC
ATTTACTAAAGAATTAGCAGTCATGATTGTGAGATCTGTTAACGGTTTAGTGTTTTCTTACCCAATTGTAGAGACTATCCACAAGGACAATATTT
GTGACTTATGTTATGCGCCTGCTAGAGTTCCGGACTCCGTTCAACTTAAGGCGAAGTTGTTGGCAGAAAATGCAATCAAATCTTTTCCCGGTTGT
GGTATATTTGGTGTGGAAATGTTCTATTTAGAAACAGGGGAATTGCTTATTAACGAAATTGCCCCAAGGCCTCACAACTCTGGACATTATACCAT
TGATGCTTGCGTCACTTCTCAATTTGAAGCTCATTTGAGATCAATATTGGATTTGCCAATGCCAAAGAATTTCACATCTTTCTCCACCATTACAA
CGAACGCCATTATGCTAAATGTTCTTGGAGACAAACATACAAAAGATAAAGAGCTAGAAACTTGCGAAAGAGCATTGGCGACTCCAGGTTCCTCA
GTGTACTTATATGGAAAAGAGTCTAGACCTAACAGAAAAGTAGGTCACATAAATATTATTGCCTCCAGTATGGCGGAATGTGAACAAAGGCTGAA
CTACATTACAGGTAGAACTGATATTCCAATCAAAATCTCTGTCGCTCAAAAGTTGGACTTGGAAGCAATGGTCAAACCATTGGTTGGAATCATCA
TGGGATCAGACTCTGACTTGCCGGTAATGTCTGCCGCATGTGCGGTTTTAAAAGATTTTGGCGTTCCATTTGAAGTGACAATAGTCTCTGCTCAT
AGAACTCCACATAGGATGTCAGCATATGCTATTTCCGCAAGCAAGCGTGGAATTAAAACAATTATCGCTGGAGCTGGTGGGGCTGCTCACTTGCC
AGGTATGGTGGCTGCAATGACACCACTTCCTGTCATCGGTGTGCCCGTAAAAGGTTCTTGTCTAGATGGAGTAGATTCTTTACATTCAATTGTGC
AAATGCCTAGAGGTGTTCCAGTAGCTACCGTCGCTATTAATAATAGTACGAACGCTGCGCTGTTGGCTGTCAGACTGCTTGGCGCTTATGATTCA
AGTTATACAACGAAAATGGAACAGTTTTTATTAAAGCAAGAAGAAGAAGTTCTTGTCAAAGCACAAAAGTTAGAAACTGTCGGTTACGAAGCTTA
TCTAGAAAACAAGTAA
```

Figure S15. Input the sequence from the *S. cerevisiae* genome to be deleted.

3. Confirm the sequence has been correctly located in the genome. Confirm with “y+enter” or abort with “n+enter” (figure S16).

```
Target sequence found:
Chromosome: chr15
Position: 564475-566191
Orientation: reverse

Proceed with deletion design? [y/N]:
```

Figure S16. Location of the target sequence for ADE2 printed to the terminal for user confirmation.

4. The script designs a deletion cassette and a set of screening primers. The expected PCR product sizes for the parent strain, initial URA3 positive transformation and after marker removal by fluoro-orotic acid (FOA) counter selection (see note 5). Confirm with “y+enter” or abort with “n+enter” (figure S17).

```
⠹ Designing deletion cassette...
⠹ Designing screening strategy...

Screening Strategy:
┌──────────────────────┬──────────────┐
│ Stage                │ Product Size │
├──────────────────────┼──────────────┤
│ Parent strain        │ 2510         │
│ After transformation │ 1887         │
│ Final deletion       │ 794          │
└──────────────────────┴──────────────┘

Screening Primers:
Forward: TGCTCCAGGTTTGATTCGTGA
Reverse: ACCTTTTGATGCGGAATTGACT

Save deletion design? [y/N]: █
```

Figure S17. Screening primers and expected PCR product sizes printed to the terminal.

5. Enter a name for the output files. The script will save a map image, *.gb*, *.fasta* and primer list to *./output*. With the default parameters provided the designed cassette will be 1713 bp, so can be readily ordered as a single piece of synthetic, linear dsDNA.
6. Transform cassette into *S. cerevisiae* using the PEG/LiOAc method as above. We typically use 500 ng of linear DNA and plate 10% of the transformation mixture on a SC-ura plate and the remaining cells on a second plate to ensure sufficient individual colonies are recovered for subsequent screening.
7. [optional]. At this stage it can be helpful to purify colonies by re-streaking on SC-ura, particularly where the deletion is expected to decrease fitness as the parent strain will be resistant to FOA, care should be taken to avoid transferring it to subsequent steps.
8. [optional] confirm integration by PCR. In this case ade2 gives an obvious pink[5] colony phenotype (figure S18). The 100% positive rate observed here is typical for this initial transformation where the mutant is viable but may be lower where the deletion has a substantial effect on cell fitness, and false positives will be observed if the deletion is lethal.

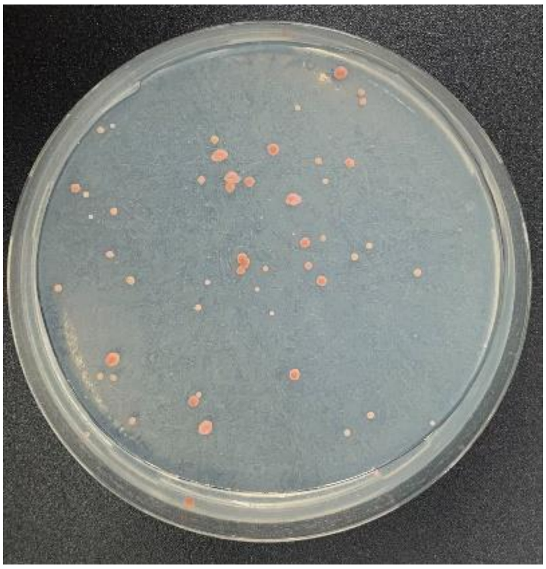

Figure S18. BY4741 ade2 deletion mutants appear pink to red.

9. Outgrow in YPD to allow for recombination between the tandem repeats to remove the marker. The rate of recombination increases with the repeat length. We optimised the default repeat during our own work over many loci, but as seen in this example, and in the initial report by Akada et al,[2] repeat lengths as short as 40 bp can work for some loci. In our experience longer repeat lengths tend to give more efficient marker removal at the cost of lower specificity in the initial transformation, meaning the DNA fragment is inserted at the incorrect locus. We have provided default parameters that work well in our hands across many genomic loci but encourage users to adjust repeat lengths first in optimising this workflow for their own work.
10. Counter select against the URA3 marker by plating 100 uL of the YPD culture on SC + 1mg/mL FOA (see note 6 for information on preparing these plates).
11. Verify marker loss by PCR. Note the PCR shown below used a set of primers flanking ADE2 already available to us, rather than those designed by PYEAST, so sizes seen here do not correspond to those shown in figure S19.

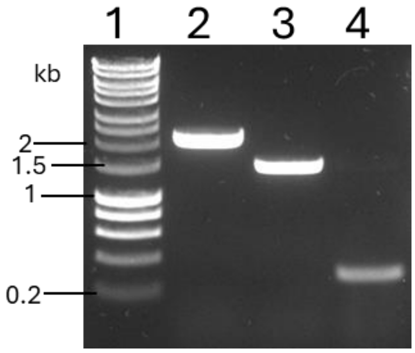


Figure S19. Gel electrophoresis confirming the ade2 transformations. Sample order: 1- Marker (hyper ladder 1kb, Bioline); 2- parent strain (ex. size 2026 bp); 3- URA3 positive strain (ex. size 1403 bp); 4- FOA resistant strain (ex, size 310 bp).

**Replacement of the promoter of the Flo1 gene using the replace command**

1. In shell program (powershell, cmd on windows, bash on linux or terminal on Mac), navigate to the directory where the git repository was cloned and run the replace command (figure S20). In this example we use the repeat_length option to set this to 40 bp (default 160).

```
PS C:\Users\loa012\Python\PYEAST> uv run pyeast replace --repeat_length 40
```

Figure S20. Running the replace command in a terminal using uv. Note the repeat_length option is used to change the size of this feature.

2. Input the deletion sequence. Note, as for the delete command, care must be taken not to copy in return keys, which appear in both *.fasta* and *.gb* files. In this example we input the sequence 500 bp immediately upstream of the Flo1 gene, a promoter which is inactive in the background strain (figure S21).

```
Enter the DNA sequence you want to replace:
Use only A, T, G, and C
Sequence: CTCATTAATTGCCCTCACAAGAATTTGGAAGTGCGTAGAACAGGTAAAAGATTGTACTACAGAGGTATTGTGGAACCTTCTACAG
TACTTCGGAATACACCTAAAAGGTTGTTGGATGCTAAATTTAGCAAAAGTCTTTTTTAGCTCACTATTAGGCTTGTTAAAGTCTGAAATTGTTGA
AAGGCACTCAAAAAGATAAATCAACAATCAGCATTAACGGCACAGTTGAAAGAGTCACCCACTTGAAATTAGCTCGGTTATCAAATATAATTATC
TCTGGTAAAGAGCTCTGCAGCAGGGTTAATCTATTCGCATACTTACGCTGTAGGAACATTTTATTATTAGGATCCGACTACTGCCTACATATTTA
TTCGGAAGGCATGATGTCGAAAATTTTTGAGCTTATAAAAGGAACATATTTCACTCTTGCTCGTTTGATGTAAGCTCTCTTCCGGGTTCTTATTT
TTAATTCTTGTCACCAGTAAACAGAACATCCAAAA
```

Figure S21. Input the sequence from the *S. cerevisiae* genome to be replaced.

3. Verify the correct location has been identified then select the component library (figure S22A) and, from the resulting list, the sequence to replace the input. Use "tab" to autocomplete selections, in this example we select the strong constitutive promoter pTef1 form the *Saccharomyces cerevisiae* library (figure S22B).

a)

```
Target sequence found:
Chromosome: chr01
Position: 202904-203404
Orientation: forward
data\component libraries
    Available Component
        Directories

 Name
 Saccharomyces cerevisiae

Select components directory (TAB for suggestions): Saccharomyces cerevisiae
```

b)

```
Select a sequence to insert:
Sequence name: pTEF1
```

Figure S22A) Verification of the correct location in the *S. cerevisiae* genome and select directory for replacement sequence. B) Select replacement sequence.

4. Select the position of the URA3 marker relative to the replacement sequence from up or down stream. In this example we select up, which will place the pTef1 promoter directly next to the Flo1 gene, causing it to be expressed in the initial URA3 transformant. Expression of Flo1 confers a flocculation phenotype, which can be readily identified in liquid culture (see figure 25B) and by colony morphology.
5. Select the position of the ura3 marker relative to the replacement sequence, by inputting either up or down and pressing "enter" (figure S23). Optionally, "tab" can be used for autocompletion.

```
Do you want the URA3 marker upstream or downstream of the replacement?
Position (up/down): up
                    up
```

Figure S23. Selection of relative orientation of the ura3 marker and replacement sequence.

6. The screening primers and the expected PCR product sizes for each stage of the process are printed to the terminal (figure S24). (See note 5)

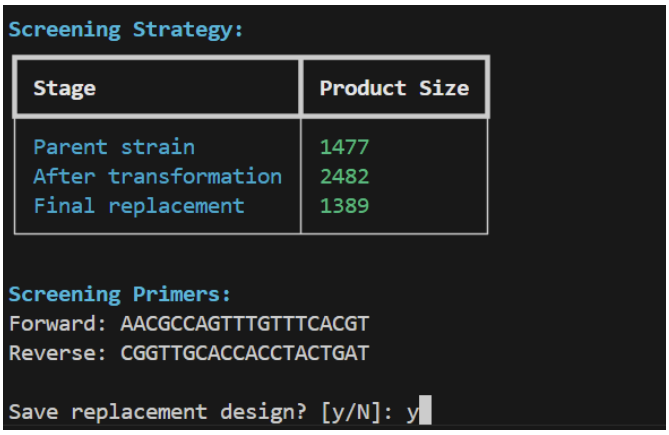


Figure S24. Screening primers and expected product sizes printed to the terminal.

7. Input a name for the output files. A sequence map, pair of screening primers, *.fasta* file and *.gb* file will be saved to the output directory. The final length of the cassette will depend on the replacement sequence selected and the parameters used. With default parameters the cassette length is 1613 bp + length of the replacement sequence. Linear synthetic dsDNA up to 3000 bp are available from most suppliers, so replacement sequences of up to 1387 bp can be input using the default parameters and remain inside this window. For longer sequence replacements we would typically create a new integration site in *.data/integration sites* and use the integrate command.
8. Transform cassette into *S. cerevisiae* using the PEG/LiOAc method as above[1]. We typically use 500 ng of linear DNA and plate 10% of the transformation mixture on a SC-ura plate and the remaining cells on a second plate to ensure sufficient individual colonies are recovered for subsequent screening.
9. [optional]. At this stage it can be helpful to purify colonies by re-streaking on SC-ura, particularly where the deletion is expected to decrease fitness.
10. [optional] Confirm integration by PCR. As with the delete command we typically observe 100% correct integrations for this initial transformation where there is no significant loss of fitness.
11. Outgrow in YPD to allow for recombination between the tandem repeats to remove the marker. The rate of recombination increases with the repeat length. The default repeat length (160 bp) has been taken from optimisation of the deletion of the Ade2 locus.
12. Counter select against the URA3 marker by plating 100 µL of the YPD culture on SC + 1 mg/mL FOA (see note 6 for information on preparing these plates).
13. Verify marker loss by PCR. Note the PCR shown below used a set of primers flanking the region upstream of the Flo1 gene, so band sizes do not correspond to those shown in figure S25A.

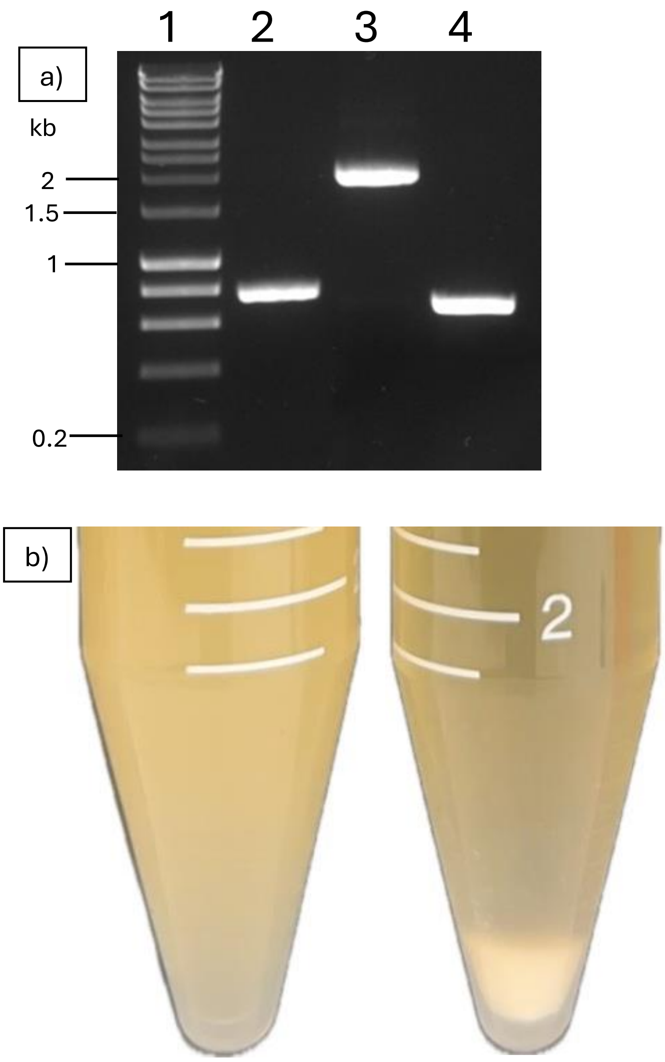


Figure S25 A) Gel electrophoresis confirming the pFLo1 replacements. Sample order: 1- Marker (hyper ladder 1kb, Bioline); 2- parent strain (ex. size 740 bp); 3- URA3 positive strain (ex. size 1745 bp bp); 4- FOA resistant strain (ex, size 652 bp). B) wild type (left) and flocculant cells (right) after replacement of the Flo1 promoter. Cells were grown in YPD overnight at 30°C, 200 rpm, briefly vortexed and allowed to stand vertically for 30 s before imagining.

Code, documentation and demonstration data from the above examples is available at https://doi.org/10.5281/zenodo.15393310.

**Notes**

1. The number of colonies generated decreases with the complexity of the assembly, as reported previously. [3,4] Plasmid assemblies tend to result in 10-100x more colonies than transformations that involve integration in the genome and so can often accommodate significantly more complex assemblies.

2. Plasmid preparations from *S. cerevisiae* contain relatively little plasmid. We recommend using the whole preparation for *E. coli* transformation and plating all of the cells on selective media at the end in two dilutions (10% of cells and remaining 90% of cells). For yeast vectors containing the high copy 2µ origin expect 40-100 *E. coli* colonies in total, and for low copy vectors containing CEN6ARS4 or similar low copy number origins of replication expect 10-20 *E. coli* colonies in total.
3. Primers and templates are matched on sequence alone, so no specific naming convention or annotations are required to find the sequences in a given design. The first four cells in the *.xlxs* file must be A1: "Plate or Box ID" B1: "Position", C1: "Sequence Name" and D1: "Sequence". A template *.xlsx* file is provided in *./data*.
4. Primers and templates should be stored in plates for liquid handling machine instructions. For the templates, names to be used should be added to the TemPlate.xlsx file in *.data/templates* at the corresponding well locations. For templates with long sequences, like genomes, the contig names should be added under a proxy name (e.g. BY4741 genome) in the genome_well_mapping.tsv file with the plate and well they are stored in and the name of all the contigs in the genome file.
5. PCR product sizes calculated during delete and replace commands are not saved to a file. The user may wish to copy and save them or note the numbers down for later reference.
6. We typically prepare SC+1mg/mL FOA plates with 0.67% w/v YNB with the recommended amount of a yeast synthetic dropout media supplement and then also add the missing component (76 mg/L uracil or histidine or 380 mg/L leucine) with 2% w/v of glucose. FOA should be dissolved in DMSO at 100 mg/ml then filter sterilized through an organic solvent compatible filter (typically PTFE or Nylon, DMSO will dissolve PVDF and PES membranes) and added to the agar immediately before pouring.

**References**

(1) Gietz, R. D.; Schiestl, R. H.; Willems, A. R.; Woods, R. A. Studies on the Transformation of Intact Yeast Cells by the LiAc/SS-DNA/PEG Procedure. *Yeast* **1995**, *11* (4), 355–360. https://doi.org/10.1002/yea.320110408.

(2) Akada, R.; Kitagawa, T.; Kaneko, S.; Toyonaga, D.; Ito, S.; Kakihara, Y.; Hoshida, H.; Morimura, S.; Kondo, A.; Kida, K. PCR-Mediated Seamless Gene Deletion and Marker Recycling in Saccharomyces Cerevisiae. *Yeast* **2006**, *23* (5), 399–405. https://doi.org/10.1002/yea.1365.

(3) Kuijpers, N. G.; Solis-Escalante, D.; Bosman, L.; van den Broek, M.; Pronk, J. T.; Daran, J.-M.; Daran-Lapujade, P. A Versatile, Efficient Strategy for Assembly of Multi-Fragment Expression Vectors in Saccharomyces Cerevisiae Using 60 Bp Synthetic Recombination Sequences. *Microbial Cell Factories* **2013**, *12* (1), 47. https://doi.org/10.1186/1475-2859-12-47.

(4) Finnigan, G. C.; Thorner, J. Complex in Vivo Ligation Using Homologous Recombination and High-Efficiency Plasmid Rescue from Saccharomyces Cerevisiae. *Bio-protocol* **2015**, *5* (13), e1521. https://doi.org/10.21769/BioProtoc.1521.
